# Vesicle Picker: A tool for efficient identification of membrane protein complexes in vesicles

**DOI:** 10.1101/2024.07.15.603622

**Authors:** Ryan Karimi, Claire E. Coupland, John L. Rubinstein

## Abstract

Electron cryomicroscopy (cryo-EM) has recently allowed determination of near-atomic resolution structures of membrane proteins and protein complexes embedded in lipid vesicles. However, particle selection from electron micrographs of these vesicles can be challenging due to the strong signal contributed from the lipid bilayer. This challenge often requires iterative and laborious particle selection workflows to generate a dataset of high-quality particle images for subsequent analysis. Here we present Vesicle Picker, an open-source program built on the Segment Anything model. Vesicle Picker enables automatic identification of vesicles in cryo-EM micrographs with high recall and precision. It then exhaustively selects all potential particle locations, either at the perimeter or uniformly over the surface of the projection of the vesicle. The program is designed to interface with cryoSPARC, which performs both upstream micrograph processing and downstream single particle image analysis. We demonstrate Vesicle Picker’s utility by determining a high-resolution map of the vacuolar-type ATPase from micrographs of native synaptic vesicles (SVs) and identifying an additional protein or protein complex in the SV membrane.

## Introduction

Membrane proteins play critical roles in cells, including in signalling, transmembrane transport, maintenance of ion gradients, and ATP synthesis. Single particle electron cryomicroscopy (cryo-EM) has emerged as an indispensable tool for the structural study of membrane proteins. This technology has revealed the atomic structures of biologically important protein complexes, such as ATP synthases, respiratory complexes and super-complexes, and a variety of G-protein coupled receptors (Guo *et al*, 2017; Vinothkumar *et al*, 2014; Zhu *et al*, 2016; Letts *et al*, 2016; Liang *et al*, 2017; Zhang *et al*, 2017). Most cryo-EM studies of membrane proteins employ detergents to extract an endogenous or over-expressed protein of interest from a membrane before purification. These detergents can disrupt delicate protein-protein and protein-lipid interactions, both of which may be important for protein function (Yin & Flynn, 2016; Virji & Knowles, 1978). Several reagents have been developed for cryo-EM imaging of membrane proteins without detergents, such as amphipols (Tribet *et al*, 1996), saposins (Frauenfeld *et al*, 2016), lipid nanodiscs enclosed by a membrane scaffolding protein (Borch & Hamann, 2009), and peptidiscs (Carlson *et al*, 2018). However, it is still necessary to extract the protein from the membrane with detergent prior to exchange into these detergent-free systems. Styrene-maleic acid block co-polymer lipid particles (SMALPs) allow extraction of membrane proteins directly from lipid bilayers without detergents (Knowles *et al*, 2009) but only a small number of protein structures have been determined with this method (Sun *et al*, 2018; Qiu *et al*, 2018; Yu *et al*, 2021; Flegler *et al*, 2020; Tascón *et al*, 2020; Yoder & Gouaux, 2020). Amphiphilic peptides have also been used to extract integral membrane proteins from endogenous lipid bilayers (Yeh *et al*, 2005), but to our knowledge they have not been used for high resolution cryo-EM to date.

Although there are still few examples, and most are quite recent, cryo-EM has also been used to determine structures of membrane proteins embedded within lipid vesicles. Specimens for these experiments have been prepared in three different ways. First, detergent-solubilized and purified membrane proteins have been reconstituted into liposomes (Wang & Sigworth, 2009; Yao *et al*, 2020; Lee & MacKinnon, 2018; Grba *et al*, 2024), which has allowed study of membrane proteins in the presence of a transmembrane ion gradient. Second, cell-derived lipid vesicles have been imaged following over-expression of a protein of interest, with the further advantage of potentially preserving some protein-lipid and protein-protein interactions. Finally, native membranes have been imaged directly, either synaptic vesicles (SVs) isolated from mammalian brain (Coupland *et al*, 2024; Wang *et al*, 2024) or mitochondria deposited directly on a cryo-EM grid (Zheng *et al*, 2024). This third approach has allowed observation of fully native protein-protein and protein-lipid interactions. Native vesicles have also allowed cryo-EM structure determination for an endogenous membrane-associated protein complex (Fu & MacKinnon, 2024).

Using cryo-EM to determine high-resolution structures of proteins in vesicles involves image processing challenges that do not exist in standard cryo-EM workflows. The first task in cryo-EM image analysis is the identification of an initial set of particle images, which is often initiated through a combination of manual selection of exemplar images and template matching, either with a high signal-to-noise ratio (SNR) reference image or a geometric shape. When imaging lipid vesicles, two problems emerge. First, the signal from the lipid bilayer is much stronger than the signal from the protein of interest. Consequently, the lipid bilayer may partially or fully obscure the proteins within, leading to difficulty in identifying bilayer-embedded particles.

Second, signal from the bilayer can dominate image alignment and 2D classification, preventing coherent averaging of the protein and calculation of class averages that show the particle of interest. Despite these complications, manual selection of hundreds to thousands of particle images, which can be laborious, may allow training of the particle selection algorithm Topaz (Bepler *et al*, 2019), leading to a reasonable starting set of particle images (Coupland *et al*, 2024; Grba *et al*, 2024; Wang *et al*, 2024; Zheng *et al*, 2024). However, when selecting particles in this way it can be challenging to distinguish proteins embedded in the membrane of intact vesicles, which can maintain an ion gradient, from proteins embedded in disrupted vesicles or patches of lipid bilayer.

To select particles that are embedded in the membrane of intact vesicles, it is first necessary to identify these vesicles in micrographs. Previous work demonstrated that the centers and radii of vesicles in micrographs can be determined with template matching using synthetic vesicle images (Wang *et al*, 2006; Liu & Sigworth, 2014). This approach requires manual intervention in the initial selection of vesicles in micrographs for template matching and likely performs best for spherical vesicles imaged with a high SNR. Past application of the method was facilitated by acquisition of defocus pairs of images to enhance the SNR, slowing down micrograph acquisition and processing.

Here, we present Vesicle Picker, a program built on the Segment Anything (SA) model (Kirillov *et al*, 2023). The program allows unsupervised identification of vesicles in cryo-EM micrographs and selection of potential particle positions, either at the vesicle perimeter in images or distributed on the vesicle surface. The program achieves high recall and precision when benchmarked against a dataset of manually annotated vesicles, and is easily included in existing workflows by interfacing with the cryoSPARC software package (Punjani *et al*, 2017) using the cryosparc-tools library. We demonstrate the utility of Vesicle Picker by identifying and analyzing a dataset of micrographs of SVs, yielding a high-resolution map of the vacuolar-type ATPase (V-ATPase) in its native membrane. The analysis also produces 2D class average images of an additional, unidentified, protein or protein complex embedded in the SV membrane.

## Methods and Results

### Computational approach and parameter optimization

SA is a recently developed foundation model for object annotation in images (Kirillov *et al*, 2023). It consists of an image encoder, a prompt encoder, and a mask decoder. The image encoder converts a grayscale or colour image to an abstract representation. The mask decoder takes the image representation and a representation of a prompt, such as a point or a box that bounds a portion of the image, and returns a binary mask. This mask indicates a continuous cluster of pixels in the input image corresponding to a predicted object, either under the point or within the bounding box provided as a prompt. SA also returns a score that describes the model’s confidence in the object indicated by the mask. By providing the image encoder with a micrograph and the prompt encoder with an evenly spaced grid of points, one can perform a search for all objects in a micrograph.

We tested the ability of SA to identify vesicles in a previously published dataset of micrographs of rat brain SVs on graphene oxide coated cryo-EM grids (Coupland *et al*, 2024) (**Fig. 1A**, *left*). Following some simple micrograph pre-processing, described in more detail below, SA readily identifies vesicles in motion corrected cryo-EM micrographs (**Fig. 1A**, *middle*). The locations of these vesicles can be used to determine potential particle locations, which may be exported to cryoSPARC or other image processing software for further analysis (**Fig. 1A**, *right)*.

**Figure 1.**
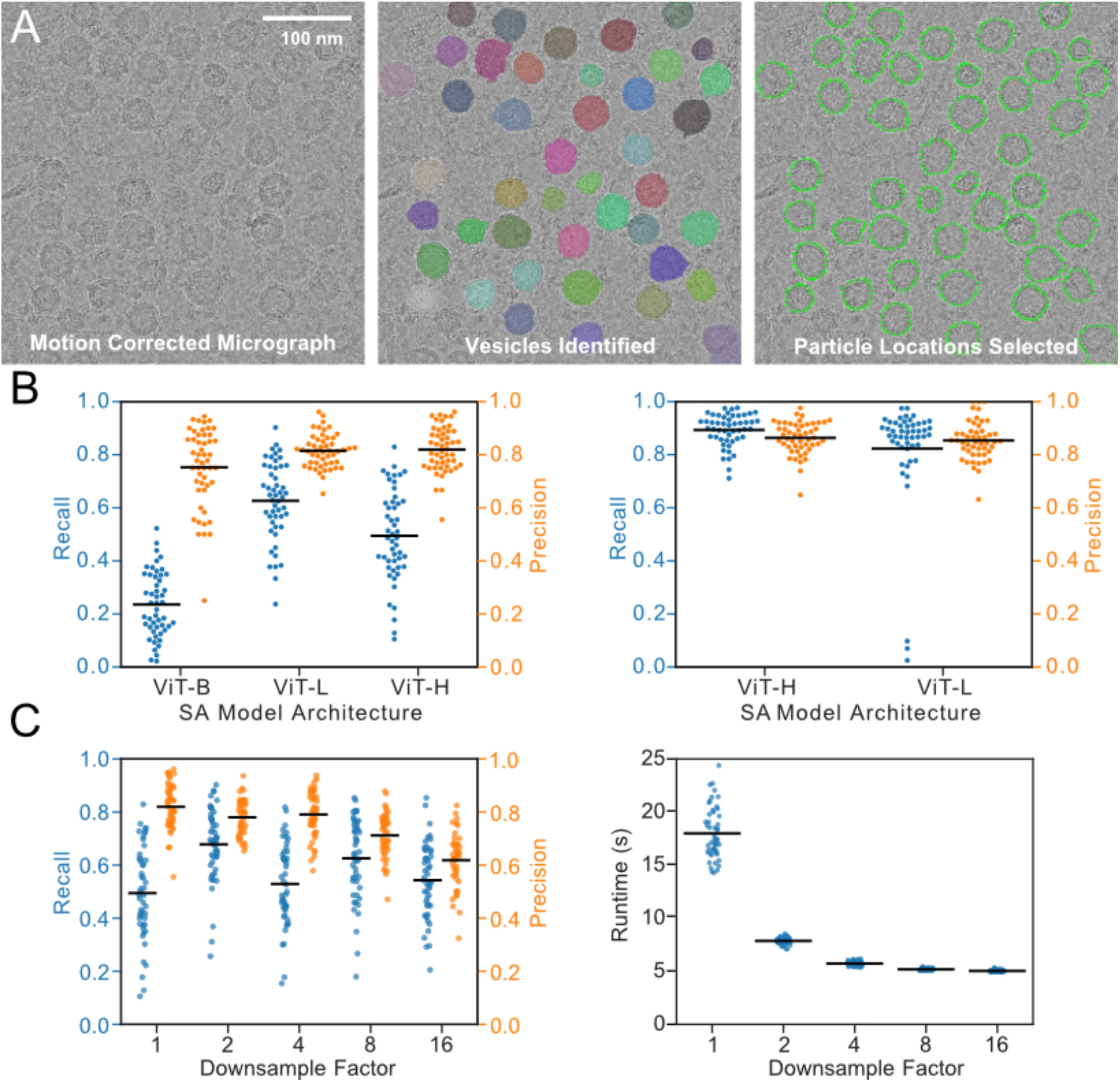
Performance of Vesicle Picker in SV identification. **A**, An example micrograph of SVs showing the input to the program (*left*), the objects identified by SA with a different colour for each object (*middle*), and the output of Vesicle Picker where potential particle locations are annotated (*right*). **B**, Unoptimized (*left*) and optimized (*right*) precision and recall with the three different Segment Anything model architectures. Each dot represents the precision or recall from one of 50 manually annotated micrographs. **C**, Precision and recall of Vesicle Picker on downsampled micrographs (*left*) and effect of downsampling on Vesicle Picker runtime (*right*).

We found that SA operating on raw vesicle micrographs often failed to identify vesicles that are apparent to a microscopist, performing with high precision (a large fraction of predicted vesicles are true vesicles) but poor recall (a small fraction of true vesicles are identified). Therefore, we sought to increase the performance of SA with additional image processing steps. These steps included pre-processing of micrographs with filters as well as application of *post hoc* selection criteria that could distinguish vesicles from non-vesicles. To quantify the effects of these additional steps, and to optimize the choice of parameters used with each, we benchmarked the algorithm’s ability to identify vesicles in a small, annotated dataset of SV micrographs. This dataset comprised 50 micrographs with boxes manually drawn around ∼2000 SVs using the LISA tool of VGG Image Annotator (Dutta & Zisserman, 2019), thereby allowing assessment of the algorithm’s precision and recall. We defined successful prediction by the model as the case where the intersection over union (IOU) of the bounding box for a predicted vesicle against a manually drawn bounding box for a true vesicle exceeded 0.5, a commonly used threshold in computer vision (Rezatofighi *et al*, 2019). We then chose parameters for micrograph pre-processing and candidate vesicle selection that provided a good balance of recall, precision, and speed in vesicle identification.

### Optimization of model architecture, micrograph downsampling, and micrograph filtering

We tested the effect of different SA model architectures on finding vesicles in the absence of pre-processing and candidate vesicle selection. Three model architectures are available for SA with different neural network sizes: Vision Transformer Base (ViT-B), Vision Transformer Large (ViT-L), and Vision Transformer Huge (ViT-H). We found that the model architectures with more parameters (ViT-H and ViT-L) outperformed the model with the fewest parameters (ViT-B) (**Fig. 1B**, *left*). While the ViT-L model performed well in identifying vesicles in raw micrographs, comparing the ViT-L and ViT-H models after optimizing pre-processing steps revealed that the ViT-H model has superior precision and recall (**Fig 1B**, *right*). We therefore used the ViT-H model. We also found that downsampling micrographs by averaging adjacent pixels prior to pre-processing and application of SA preserved precision and recall (**Fig. 1C**, *left*) while affording a substantial decrease in processing time (**Fig. 1C**, *right*). Therefore, all analysis was performed on micrographs downsampled 4×, decreasing processing time from ∼17 s/micrograph to ∼5.5 s/micrograph with a single Nvidia 3080Ti GPU.

We compared a Gaussian filter and an edge-preserving bilateral filter as a micrograph pre-processing step. The Gaussian filter replaces the value of each pixel in an image with a weighted average of neighbouring pixels, suppressing high-frequency information that can have a lower SNR than low-frequency information. The bilateral filter weights the contribution of each neighbouring pixel to the average using both a spatial proximity parameter (σ_*space*_) and an intensity similarity parameter (σ_*colour*_), downweighting the contribution of distant and dissimilar pixels and thereby preserving edges. While both types of filter increase recall substantially compared to application of SA to unfiltered micrographs (**Fig. 2A**), the edge-preserving bilateral filter with appropriately chosen parameters outperformed the Gaussian filter (**Fig. 2B**).

**Figure 2.**
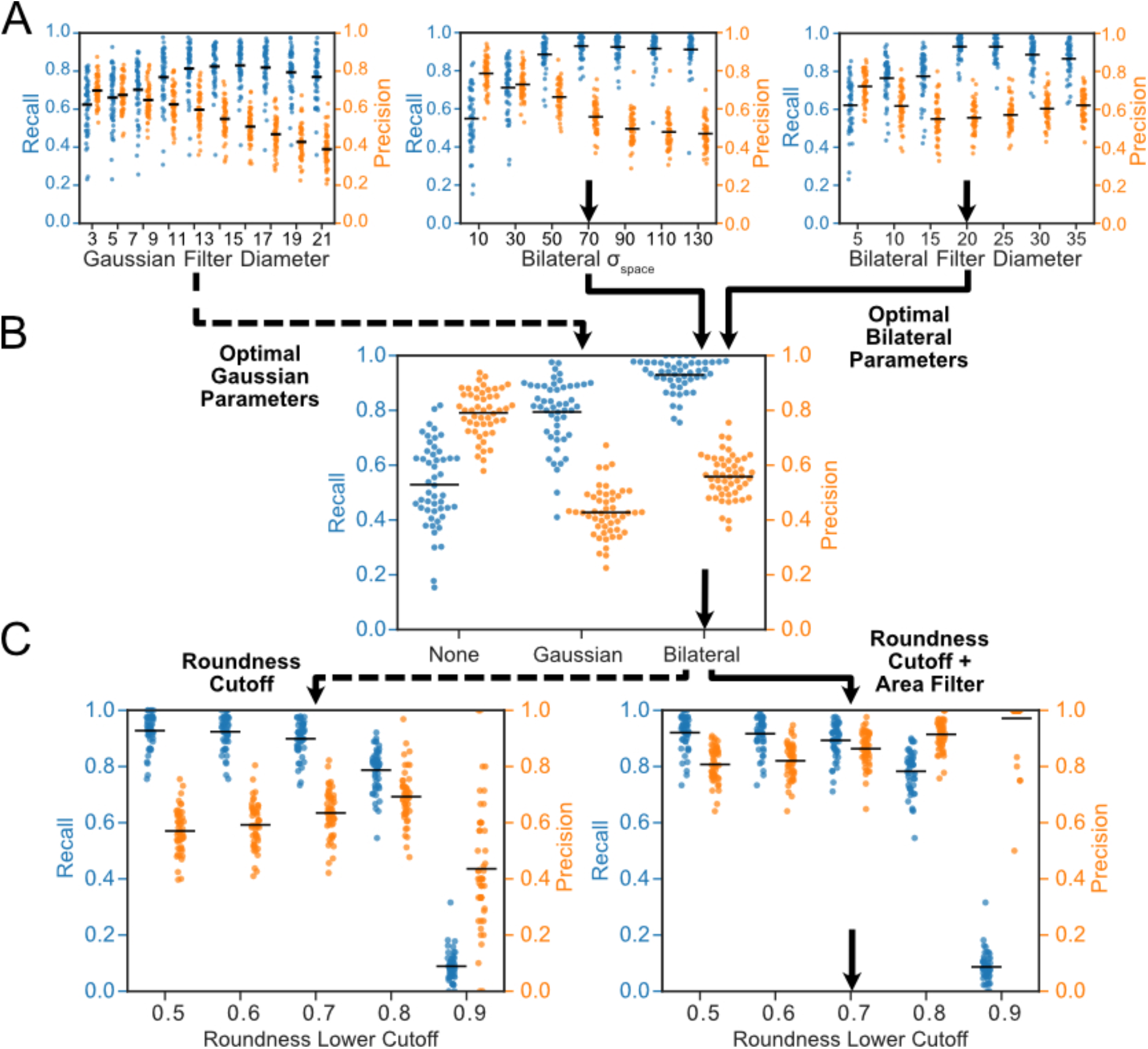
**A**, Precision and recall depending on the type and extent of lowpass filters applied to the micrograph prior to application of SA. **B**, Comparison of the precision and recall for micrographs without a filter, with an optimized Gaussian filter, and with an optimized bilateral filter. **C**, Precision and recall following application of candidate vesicle selection criteria based on vesicle roundness (*left*) and vesicle roundness with a defined maximum and minimum vesicle area between 500 and 50,000 nm^2^ (*right*).

Pre-processing filters substantially increased recall but decreased precision by causing selection of both vesicles and non-vesicles in micrographs. Therefore, we implemented *post hoc* candidate vesicle selection criteria to exclude selected objects that are not vesicles. First, we eliminated objects that were not sufficiently round. We define the roundness as 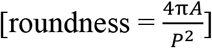 where *A* is the area of the object in the image and *P* is the length of the perimeter of the object, as computed by OpenCV (Bradski, 2000). This metric has a value of 1 for perfectly round objects and less than 1 for objects that are not round. Second, recognizing that in many experiments the expected size of the vesicles is known (e.g. SVs have an expected diameter of ∼40 nm and are consequently ∼1250 nm^2^ in projection), we enforced a minimum and maximum image area for candidate vesicle selection. Judicious choice of the roundness, minimum, and maximum area parameters allowed optimization of precision and recall for the manually-annotated dataset (**Fig. 2C**).

### Particle selection from vesicles

With a final set of vesicles identified in each micrograph, we developed an approach to convert either the area or the perimeter of each vesicle into a list of locations of potential particles. In this approach, the bounding box for each vesicle identified by SA is subdivided into patches, the size of which control the density of particle selection (**Fig. 3A**). If only side views of particles at the edges of the vesicle image are desired, the perimeter of the vesicle is found with OpenCV. The vesicle area can also be dilated before finding its perimeter to analyse particles that extend from the lipid bilayer. Assuming proteins embedded in membranes are free to rotate about a vector normal to the plane of the lipid bilayer, these side views form a single axis tomographic series and are sufficient to determine a 3D structure of the protein. Patches that do not contain any perimeter pixels are discarded. For patches that do contain perimeter pixels, the location of the perimeter pixel closest to the central pixel of the patch is recorded as the potential particle location (**Fig. 3B**, *center and right*). When particle views including top and bottom views are desired, a similar analysis is performed. However, in this case, the entire vesicle area is considered for particle selection and the location of the central pixel of each patch that belongs entirely to the vesicle image area is also recorded as a potential particle location (**Fig. 3B**, *left*). Combining the locations from patches provides a list of all potential locations at the user-controlled density (**Fig. 3C**), which can then be exported to cryoSPARC with cryosparc-tools for downstream analysis.

**Figure 3.**
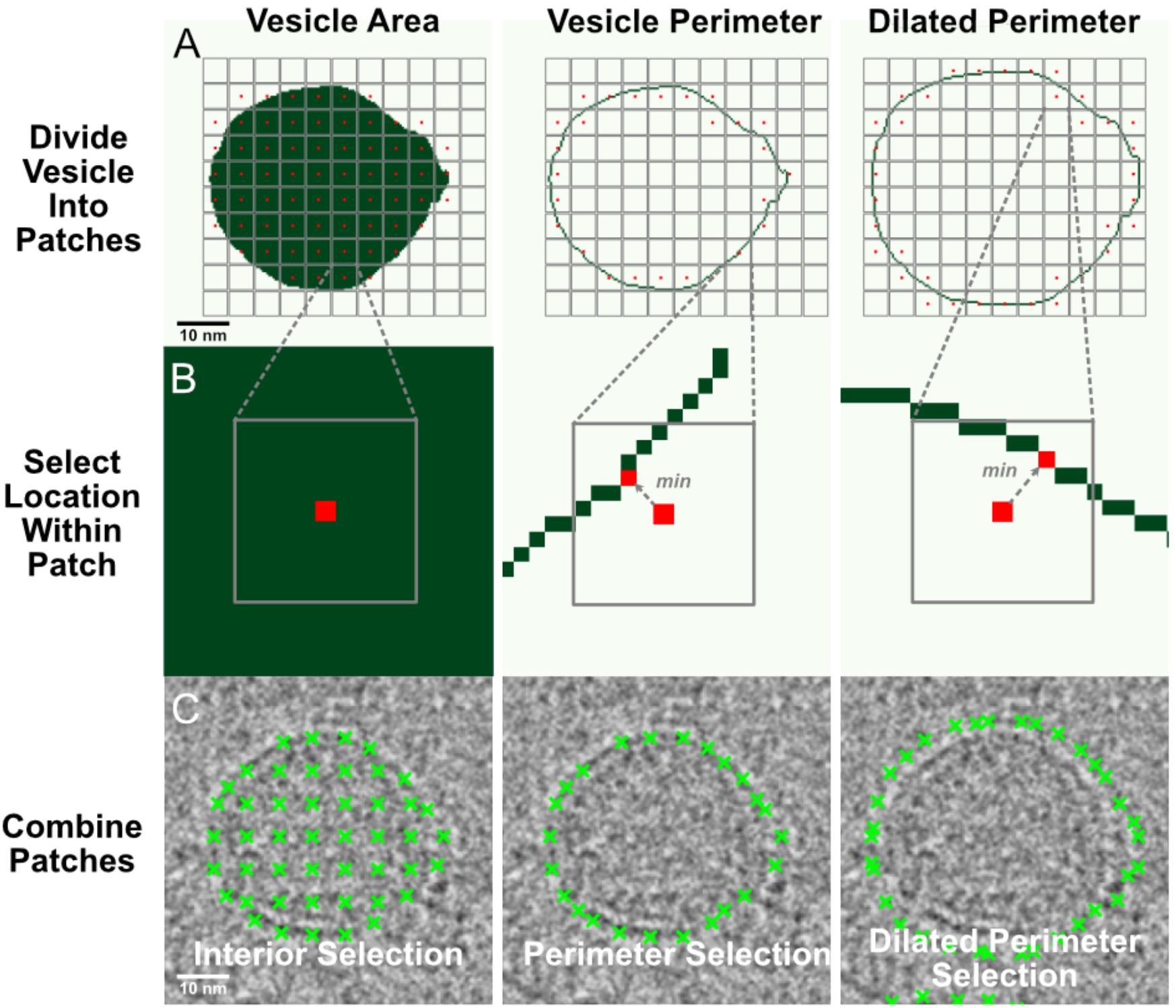
Conversion of image regions identified as vesicles to potential particle locations. **A**, An example image region identified as a vesicle (green) subdivided into patches for particle location selection. **B**, Selection of the optimal potential particle location within each patch. **C**, Potential particle locations distributed over the vesicle surface (*left*), at the vesicle image perimeter (*middle*), or offset from the vesicle image perimeter by vesicle dilation (*right*).

### Application of Vesicle Picker to V-ATPase in SV membranes

We next used the algorithm to determine a near atomic-resolution structure of the V-ATPase, a large membrane protein complex found in the membranes of many eukaryotic organelles, including SVs (Vasanthakumar & Rubinstein, 2020). We began this analysis with a dataset of ∼18,000 micrographs of SVs used previously to determine a V-ATPase structure (Coupland *et al*, 2024). All image processing was done within cryoSPARC (Punjani *et al*, 2017) or Vesicle Picker. Motion correction was performed and contrast transfer function parameters were estimated in patches prior to use of Vesicle Picker. Micrographs were transferred to Vesicle Picker with cryosparc-tools, and SVs were selected following downsampling of micrographs 4× and application of a bilateral filter with the parameters that maximized recall on the manually annotated dataset (σ_*space*_ = 71, σ_*colour*_ = 71, filter diameter = 27), as well as a minimum roundness cutoff of 0.75 and a minimum and maximum area of 500 and 50,000 nm^2^, respectively. The center of mass of the V-ATPase is offset from the bilayer edge by its large, soluble V_1_ region; consequently, we used the offset particle selection strategy described above (**Fig. 3C**, *right*, and **Fig. 4A**). A vesicle dilation of 60 Å was chosen empirically so that the selected locations corresponded with the centers of mass of the V-ATPase complexes visible in the micrographs. The density for particle selection was set to 80 Å to accommodate the ∼200 Å length of the V-ATPase complexes and enable extraction of multiple particle images per particle. These overlapping images ensure that there is at least one particle image that is well-centered for each V-ATPase complex. Particle coordinates were used in cryoSPARC to extract particle images in 392 × 392 Å boxes. These extracted particle images were subjected to multiple rounds of 2D classification and classes were selected at each iteration that showed features consistent with the V-ATPase. The resulting final 2D class average images show clear projections of the V-ATPase structure (**Fig. 4B**). Particle images in the selected 2D classes within 250 Å of another particle were removed to eliminate duplicate particle images. Particle images were next recentered using the calculated shifts from 2D classification and re-extracted in 500 × 500 Å boxes. Multi-class *ab initio* reconstruction starting with these particle images produced several maps that show the V-ATPase in its different rotational states (Zhao *et al*, 2015; Abbas *et al*, 2020). Heterogeneous refinement of the maps followed by non-uniform refinement (Punjani *et al*, 2020) allowed calculation of a map of the V-ATPase in rotational state 3 to a nominal overall resolution of 3.8 Å (**Fig. 4C and D**). This resolution is comparable to the 3.5 Å obtained with the more laborious particle selection approach used for the previous analysis of the same dataset (Coupland *et al*, 2024). The map could likely be improved further by using identified particle images to train Topaz for additional particle selection, using the calculated map to generate templates for the identification of top views by template matching, or by focused refinement of the different regions of the complex, none of which were performed for the map shown here.

**Figure 4.**
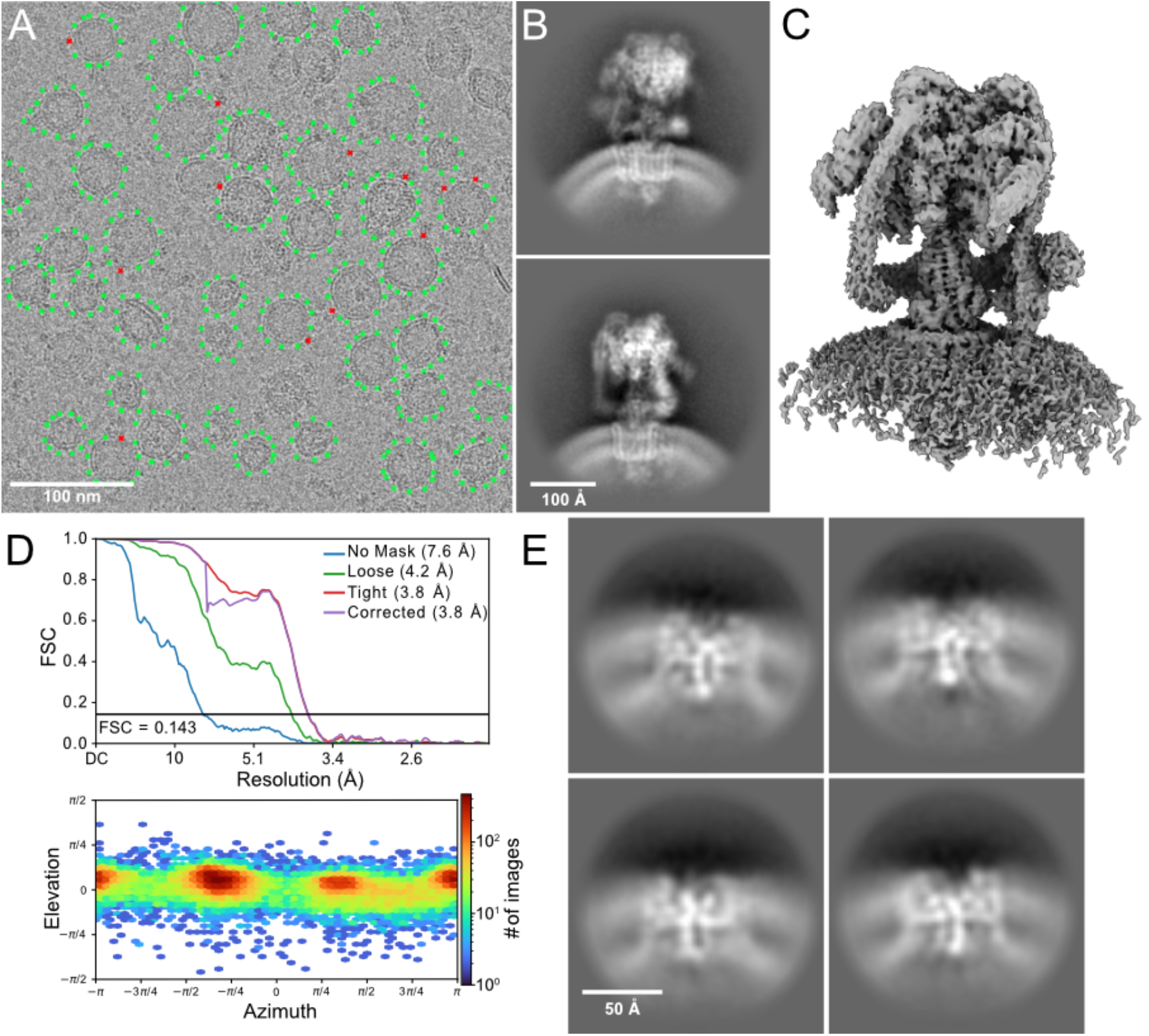
Application of Vesicle Picker to an SV dataset. **A**, An example micrograph demonstrating selection of potential particle locations offset from the vesicle image perimeter (*green crosses*) used to find V-ATPase complexes (*red crosses*) in images. **B**, 2D class average images showing the V-ATPase. **C**, A high resolution map of the V-ATPase embedded in the SV membrane. **D**, Fourier shell correlation curve (*upper*) and viewing angle distribution plot (*lower*) for the map shown in part C. **E**, 2D class average images of an unknown protein or protein complex in the SV membrane.

### Identification of an unknown protein in the SV membrane

We next used Vesicle Picker to find potential particle locations without an offset from the membrane. This procedure allowed us to identify an additional small SV protein or protein complex (**Fig. 4E**). Detection of this protein, which was not achieved in the published study of the SV micrographs, is likely the result of the focused and unbiased nature of the vesicle-constrained particle selection strategy. The ability to detect unexpected proteins in the membrane suggests that Vesicle Picker may have utility as a tool for the structural exploration of SVs and other native membranes.

## Discussion

Vesicle Picker provides an efficient, unbiased, and largely unsupervised workflow for determining structures of vesicle-embedded protein complexes. The program is underpinned by the strong performance of the SA model in identifying objects in cryo-EM micrographs. These micrographs have a much lower SNR than the images on which we presume the SA model was trained, and therefore this performance was unexpected. Recent work has shown that SA can be used in many contexts for which it was not originally designed, such as medical imaging (Zhang *et al*, 2024) and light microscopy (Archit *et al*, 2023). Unlike previous methods for defining vesicles in micrographs (Wang *et al*, 2006), Vesicle Picker does not require any prior knowledge about the morphology of vesicles present in the sample. It is therefore well-suited to process micrographs of heterogenous vesicles and native membrane vesicles containing many different proteins. It could also find application in analysis of pleomorphic virus structures.

Using the program, we determined a near atomic-resolution structure of the V-ATPase in SVs, largely recapitulating features reported recently from the same dataset (Coupland *et al*, 2024) but using a substantially more streamlined workflow. The marginally worse resolution in the map presented here (3.8 Å versus 3.5 Å previously) is likely because the number of particle images used was approximately one third of the number used for the earlier work. The smaller number of particles is likely the result of missing top views, V-ATPases in disrupted vesicles that were not selected by Vesicle Picker, and imperfect recall in identifying vesicles. This finding suggests that while the approach presented here can find particles embedded in the membranes of intact vesicles, using an approach like Topaz can still provide more particle images. If one wishes to maximize the number of particle images available for 3D reconstruction, the highest quality particle images from Vesicle Picker can be used to seed a Topaz model or for template matching to find additional views of the complex.

Finally, while Vesicle Picker helps address bottlenecks associated with particle selection in membranes, determining high resolution structures of membrane-embedded protein complexes still suffers from additional problems. In particular, generation and refinement of 3D maps would benefit from methods to compensate for the contribution of the lipid bilayer to image alignment and classification.

## Code availability

Vesicle Picker is implemented as an open-source Python package and can be installed from GitHub at https://github.com/r-karimi/vesicle-picker.

## Acknowledgments

We thank Yingke Liang, Gautier Courbon, and Justin Di Trani for helpful discussions about processing datasets of protein complexes in membrane vesicles. RK and CEC were supported by ResTraComp fellowships from the Hospital for Sick Children and RK was supported by a Canada Graduate Scholarship from the Canadian Institutes for Health Research. JLR was supported by the Canada Research Chairs program. This research was supported by Natural Sciences and Engineering Research Council grant RGPIN-2023-04676 to JLR. Cryo-EM data were collected at the Toronto High-Resolution High-Throughput cryo-EM facility, which is supported by the Canada Foundation for Innovation and Ontario Research Fund.

## Author contributions

RK found that Segment Anything could identify vesicles in micrographs and developed the software presented in this study. RK applied the software to the previously published dataset of SV images collected by CEC with insight and guidance from CEC. JLR conceived the project and supervised this study. RK and JLR wrote the paper and prepared the figures.

